## Supplementary Figures 1-13, Tables 1-6 for "C-C chemokine receptor 4 deficiency exacerbates early atherosclerosis in mice"

**Content**
Supplementary Figures 1-13
Supplementary Tables 1-6

### Supplementary Figures

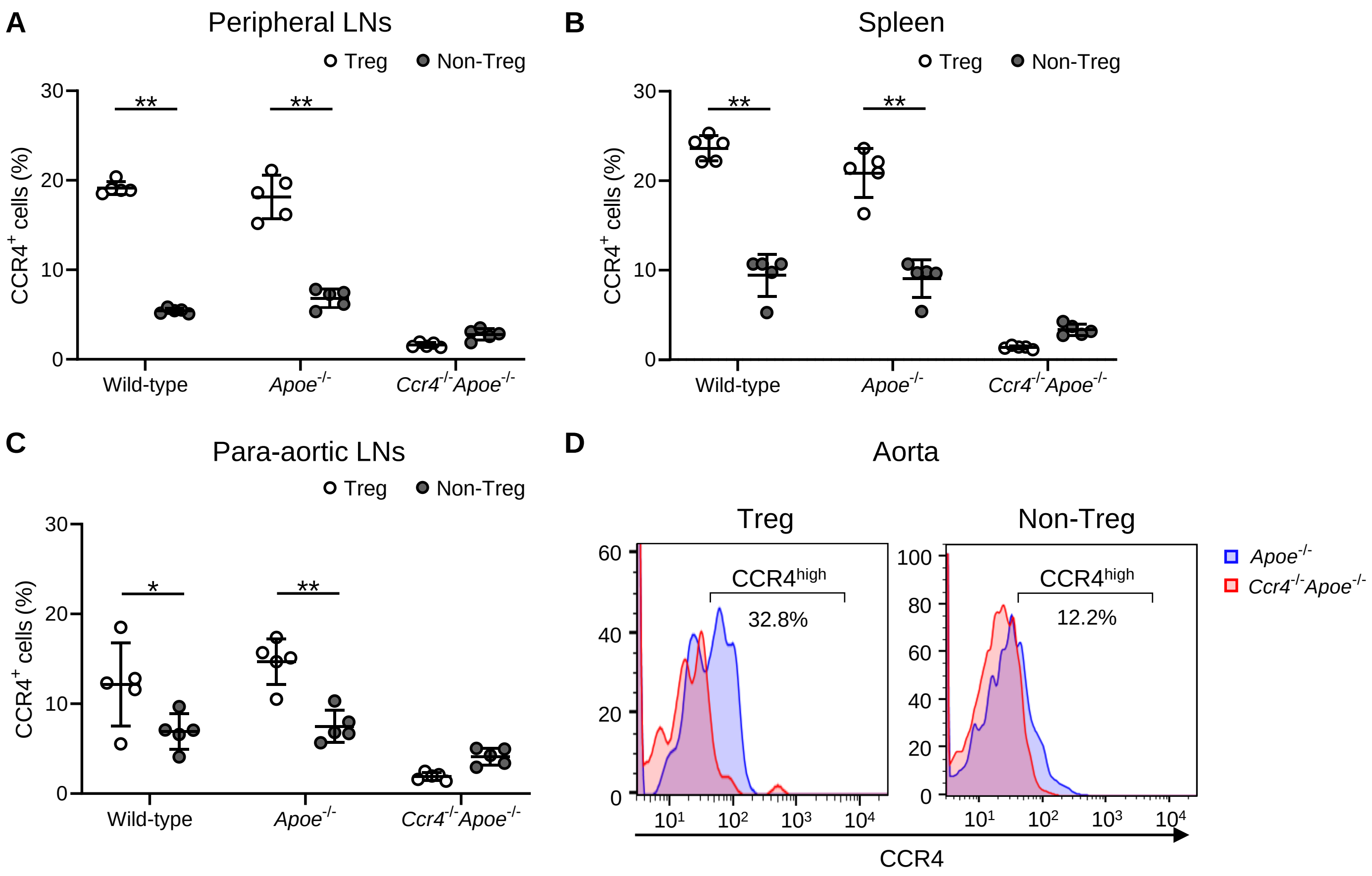


**Supplementary Figure 1. CCR4 is predominantly expressed on CD4⁺Foxp3⁺ Tregs.**
**A**-**C**, The graphs represent the proportions of CCR4⁺ cells in CD4⁺Foxp3⁺ Tregs and CD4⁺Foxp3⁻ non-Tregs in the peripheral LNs (**A**), spleen (**B**), and para-aortic LNs (**C**) of 18-week-old wild-type, *Apoe*$^{-/-}$, or *Ccr4*$^{-/-}$*Apoe*$^{-/-}$ mice assessed by flow cytometry. n=5 per group. Data points represent individual animals. Horizontal bars represent means. Error bars indicate s.d. **D**, Representative flow cytometric analysis of CCR4 expression in aortic CD4⁺Foxp3⁺ Tregs and CD4⁺Foxp3⁻ non-Tregs from 18-week-old *Apoe*$^{-/-}$ or *Ccr4*$^{-/-}$*Apoe*$^{-/-}$ mice. Pooled aortic lymphoid cells from 7 to 8 mice in each group were used. Data are representative of 2 independent experiments. **P*<0.05, ***P*<0.01; 2-way ANOVA followed by Tukey's multiple comparisons test: **A**, **B** and **C.**

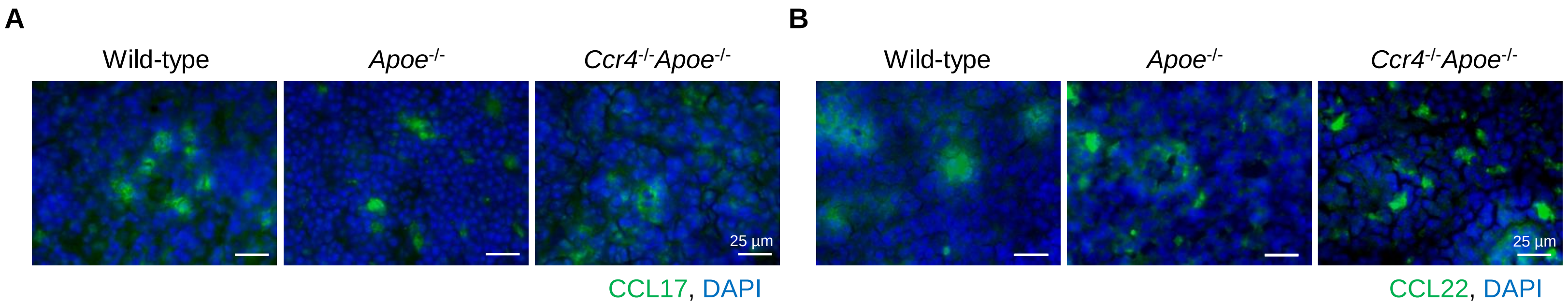


**Supplementary Figure 2. CCL17 and CCL22 are detected in peripheral LNs.**
Immunostaining for CCL17 (green) (**A**) and CCL22 (green) (**B**) in the peripheral LNs of 18-week-old wild-type, *Apoe*$^{-/-}$, or *Ccr4*$^{-/-}$*Apoe*$^{-/-}$ mice. Nuclei were stained with DAPI (blue). Data are representative of 5 mice analyzed in each group. White bars represent 25 μm as described.

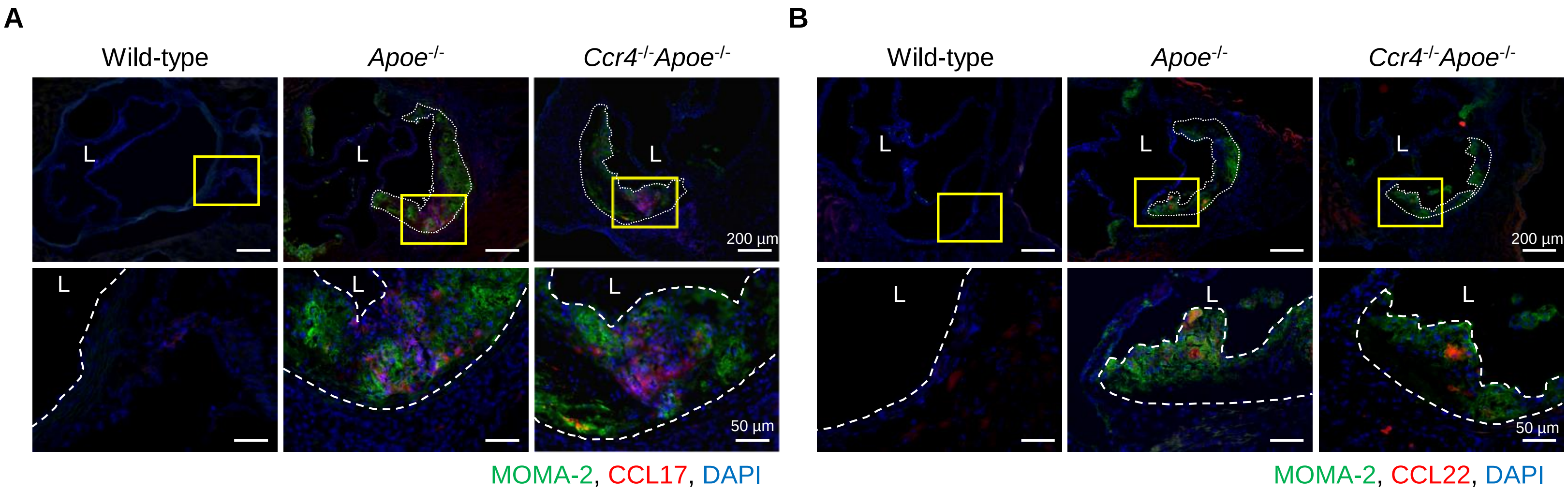


**Supplementary Figure 3. CCL17 and CCL22 are detected in atherosclerotic lesions.**
Immunostaining for for CCL17 (red) and MOMA-2 (green) (**A**) and for CCL22 (red) and MOMA-2 (green) (**B**) in the aortic sinus of 18-week-old wild-type, *Apoe*$^{-/-}$, or *Ccr4*$^{-/-}$*Apoe*$^{-/-}$ mice. Boxed area is expanded to show high-power fields. Nuclei were stained with DAPI (blue). Dashed lines demarcate atherosclerotic lesions or indicate the inner lining of arteries; L, lumen. Data are representative of 5 mice analyzed in each group. White bars represent 50 or 200 μm as described.

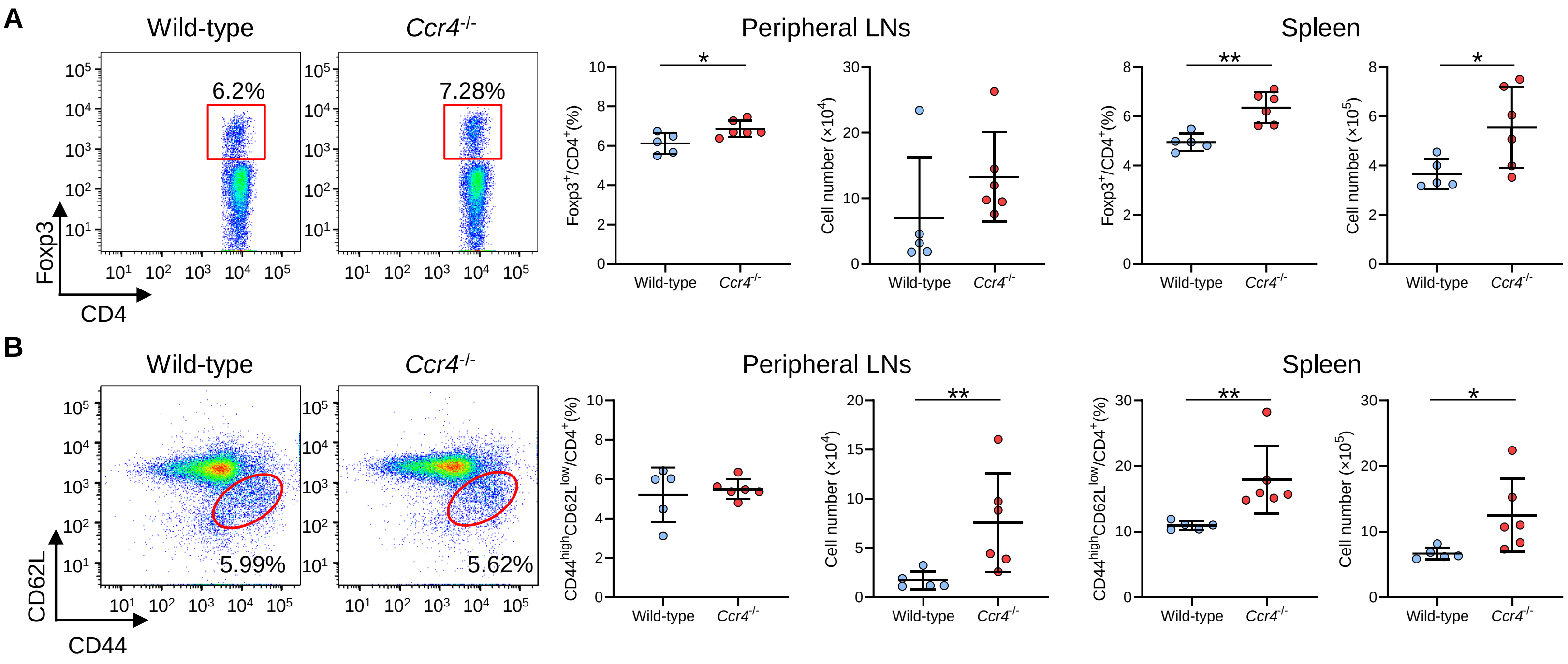


**Supplemental Figure 4. CCR4 deficiency expands peripheral Tregs and effector memory T cells in the peripheral lymphoid tissues of normocholesterolemic mice.**

Representative flow cytometric analysis of CD4$^+$Foxp3$^+$ Tregs (**A**) and CD4$^+$CD44$^{high}$CD62L$^{low}$ effector memory T cells (**B**) in the peripheral LNs and spleen of 8-week-old wild-type or *Ccr4$^{-/-}$* mice. The graphs represent the total numbers and proportions of CD4$^+$Foxp3$^+$ Tregs (**A**) and CD4$^+$CD44$^{high}$CD62L$^{low}$ effector memory T cells (**B**) in the peripheral LNs and spleen. n=5 to 6 per group. Data points represent individual animals. Horizontal bars represent means. Error bars indicate s.d. *$P$<0.05, **$P$<0.01; Mann-Whitney *U*-test: **B** second and third from the left; 2-tailed Student's *t*-test: **A** and **B** fourth from the left.

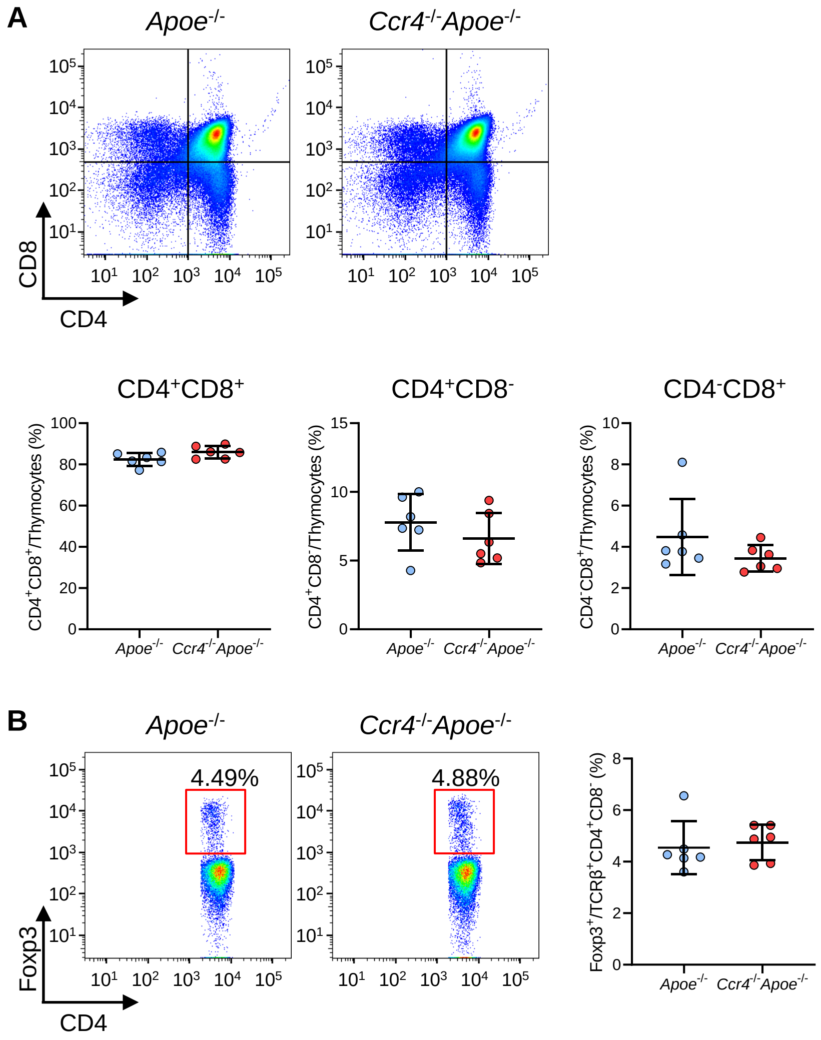


**Supplementary Figure 5. CCR4 deficiency does not affect T cell development in the thymus.**

Lymphoid cells from the thymus of 4-week-old *Apoe*$^{-/-}$ or *Ccr4*$^{-/-}$*Apoe*$^{-/-}$ mice were prepared. **A**, Representative flow cytometric analysis of CD4 and CD8 expression in thymocytes. The graphs represent the numbers and proportions of CD4/CD8 double positive (CD4$^{+}$CD8$^{+}$), CD4 single positive (CD4$^{+}$), and CD8 single positive (CD8$^{+}$) T cells among thymocytes. **B**, Representative flow cytometric analysis of Foxp3 expression among the TCR-β$^{+}$CD4$^{+}$CD8$^{-}$ population. The graph represents the proportion of Foxp3$^{+}$ Tregs among the TCR-β$^{+}$CD4$^{+}$CD8$^{-}$ population. n=6 per group. Data points represent individual animals. Horizontal bars represent means. Error bars indicate s.d.

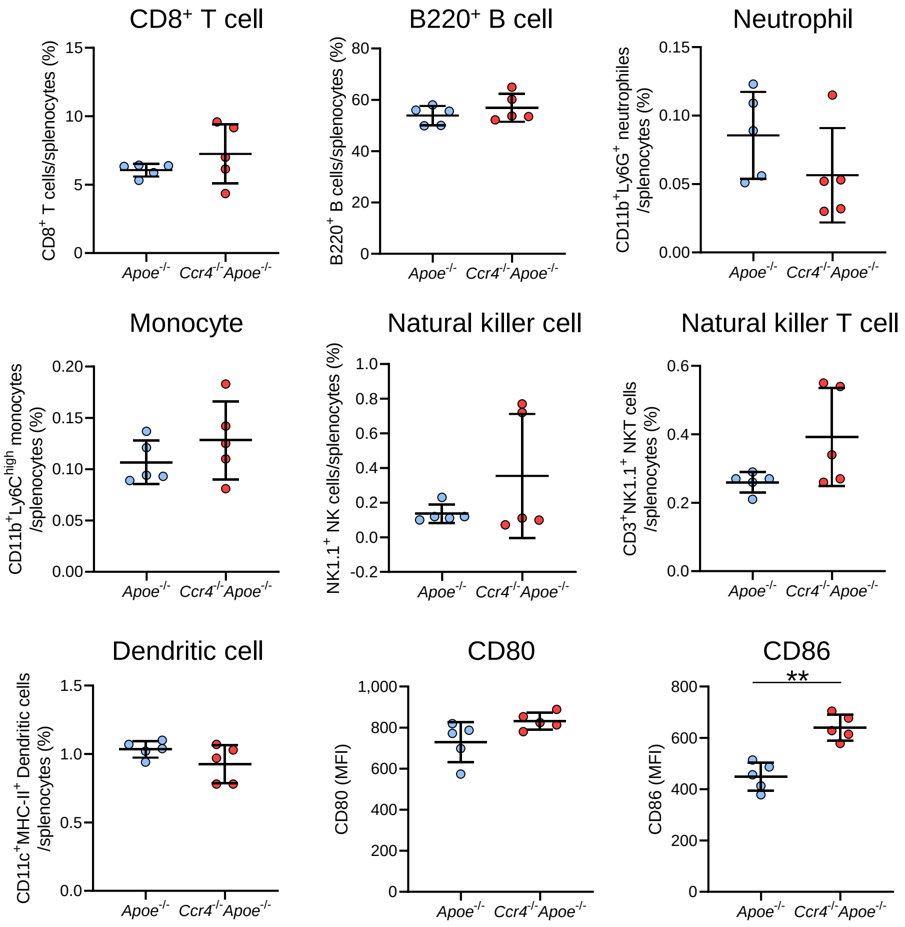


**Supplementary Figure 6. CCR4 deficiency has a minor effect on the proportions of other immune cells in spleen.**

Proportions of splenic $CD8^+$ T cells, $B220^+$ B cells, $CD11b^+Ly6G^+$ neutrophils, $CD11b^+Ly6C^{high}$ monocyte, NK cells, NKT cells, and $CD11c^+MHC\text{-}II^+$ DCs, and the expression of CD80 and CD86 on $CD11c^+MHC\text{-}II^+$ DCs in 8-week-old *Apoe*$^{-/-}$ or *Ccr4*$^{-/-}$*Apoe*$^{-/-}$ mice were determined by flow cytometry. n=5 per group. Data are representative of 2 independent experiments. Data points represent individual animals. Horizontal bars represent means. Error bars indicate s.d. $^{**}P<0.01$; 2-tailed Student's *t*-test. MFI indicates mean fluorescence intensity.

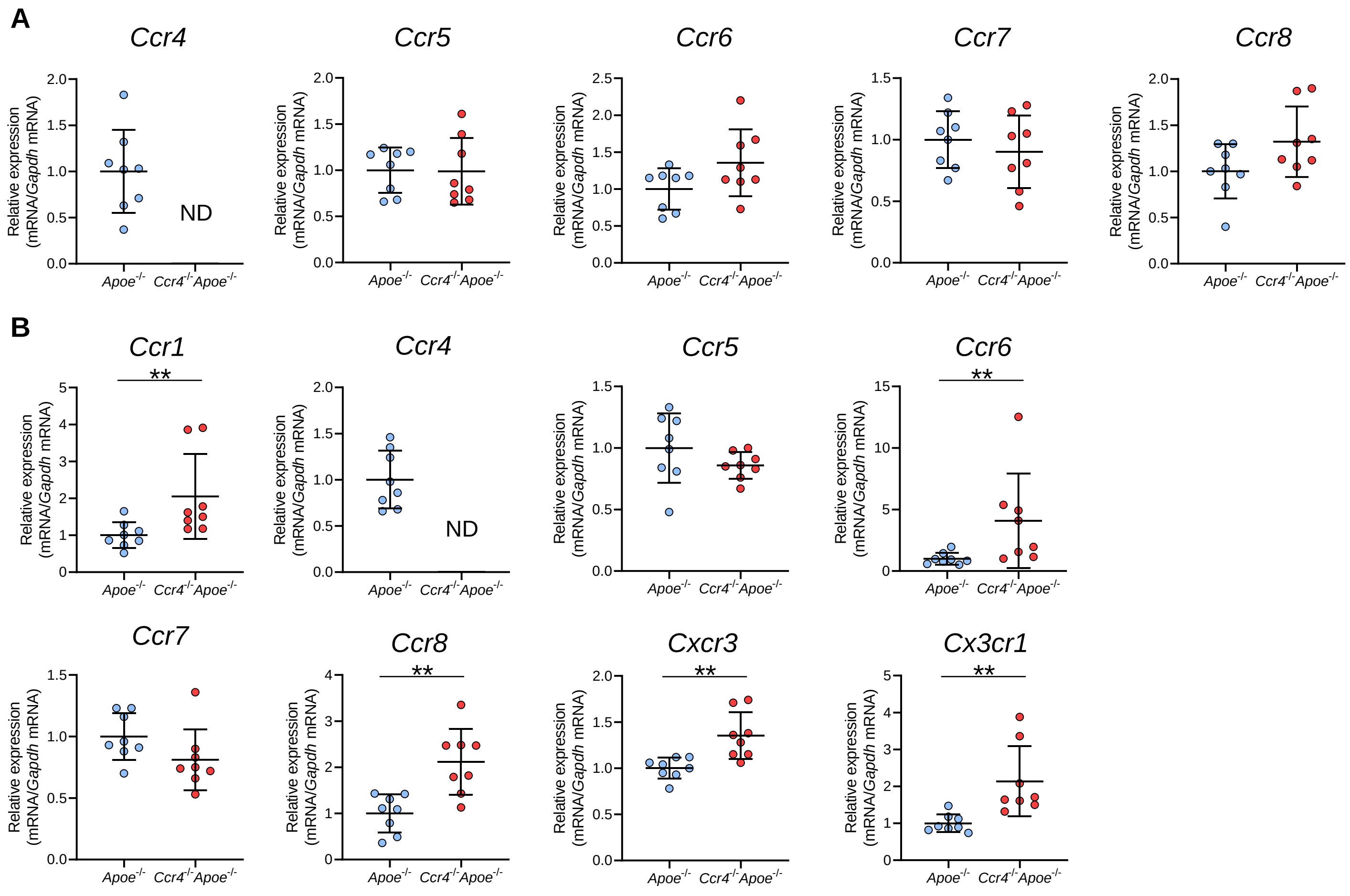


**Supplementary Figure 7. The effect of CCR4 deficiency on the expression of various chemokine receptors in splenic Tregs and non-Tregs.**

mRNA expression of chemokine receptors in splenic Tregs (**A**) and non-Tregs (**B**) from 8-week-old *Apoe*$^{-/-}$ or *Ccr4*$^{-/-}$*Apoe*$^{-/-}$ mice. The expression levels of the target genes were normalized so that the mean values in *Apoe*$^{-/-}$ mice were set to 1. n=8 per group. Data points represent individual animals. Horizontal bars represent means. Error bars indicate s.d. $^{**}P<0.01$; Mann-Whitney *U*-test: **B** top first and third from the left and bottom third from the left; 2-tailed Student's *t*-test: **B** bottom first and second from the left. ND indicates not detected.

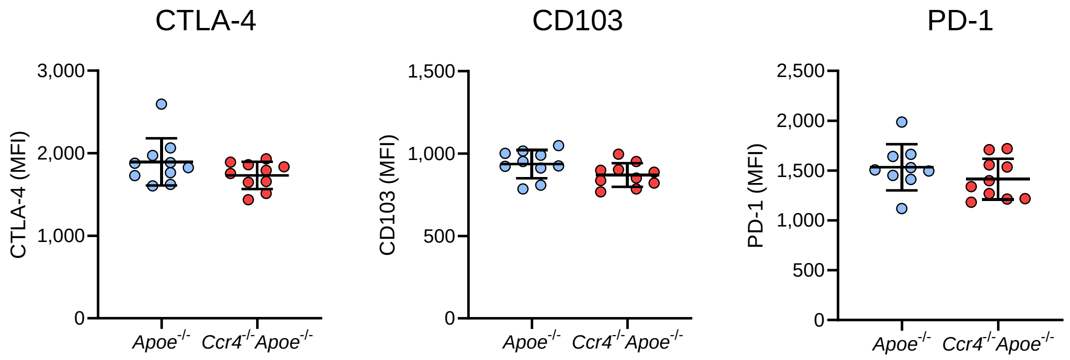


**Supplementary Figure 8. CCR4 deficiency does not affect the expression of activation- or function-associated molecules in Tregs in para-aortic LNs.**

Expression levels of CTLA-4, CD103, and PD-1 were analyzed by gating on CD4$^{+}$Foxp3$^{+}$ Tregs in the para-aortic LNs of 18- week-old *Apoe*$^{-/-}$ or *Ccr4*$^{-/-}$*Apoe*$^{-/-}$ mice. n=9 to 10 per group. Data points represent individual animals. Horizontal bars represent means. Error bars indicate s.d. MFI indicates mean fluorescence intensity.

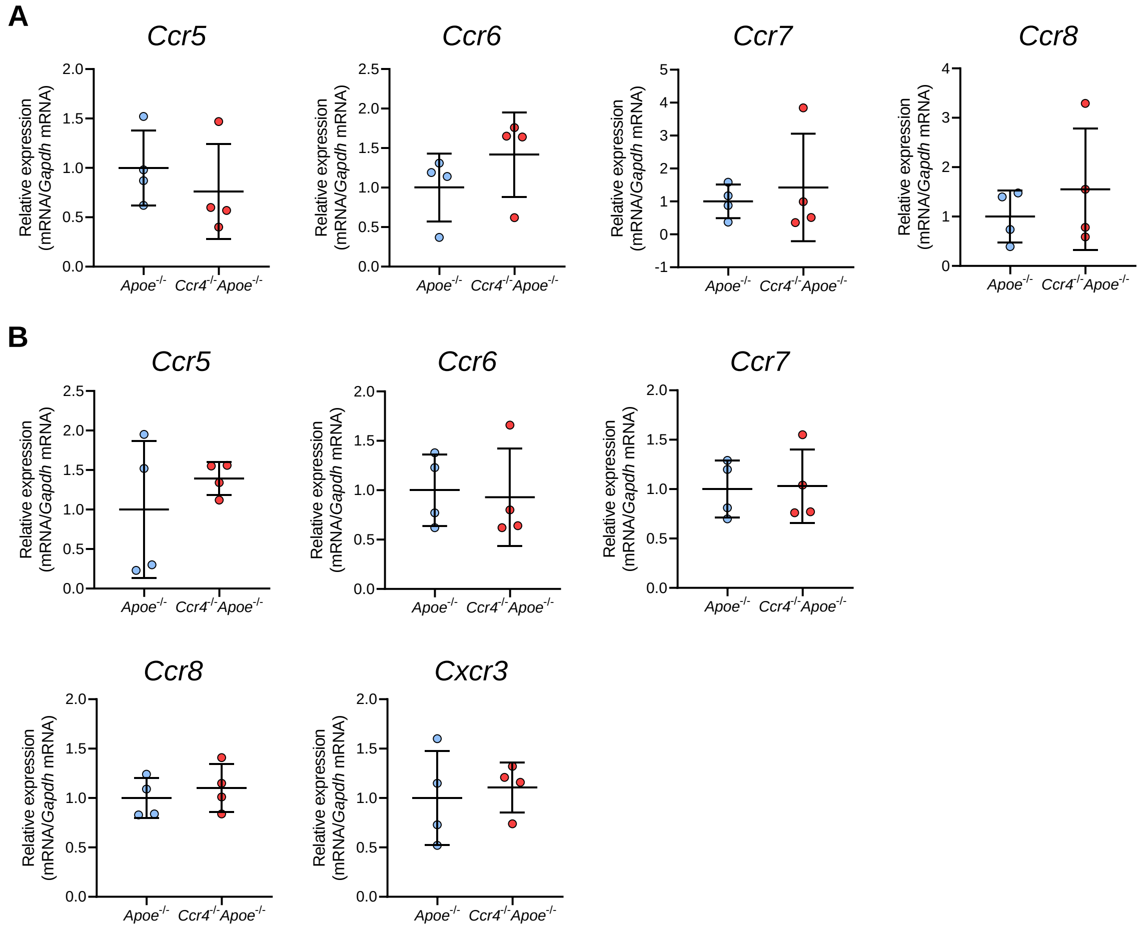


**Supplementary Figure 9. The effect of CCR4 deficiency on the expression of various chemokine receptors in Tregs and non-Tregs in para-aortic LNs.**

mRNA expression of chemokine receptors in Tregs (**A**) and non-Tregs (**B**) in the para-aortic LNs of 18-week-old *Apoe*$^{-/-}$ or *Ccr4*$^{-/-}$*Apoe*$^{-/-}$ mice. The expression levels of the target genes were normalized so that the mean values in *Apoe*$^{-/-}$ mice were set to 1. Tregs or non-Tregs purified from pooled para-aortic LNs of 9 to 10 mice were analyzed as a sample. n=4 per group. Data points represent individual pooled samples. Horizontal bars represent means. Error bars indicate s.d.

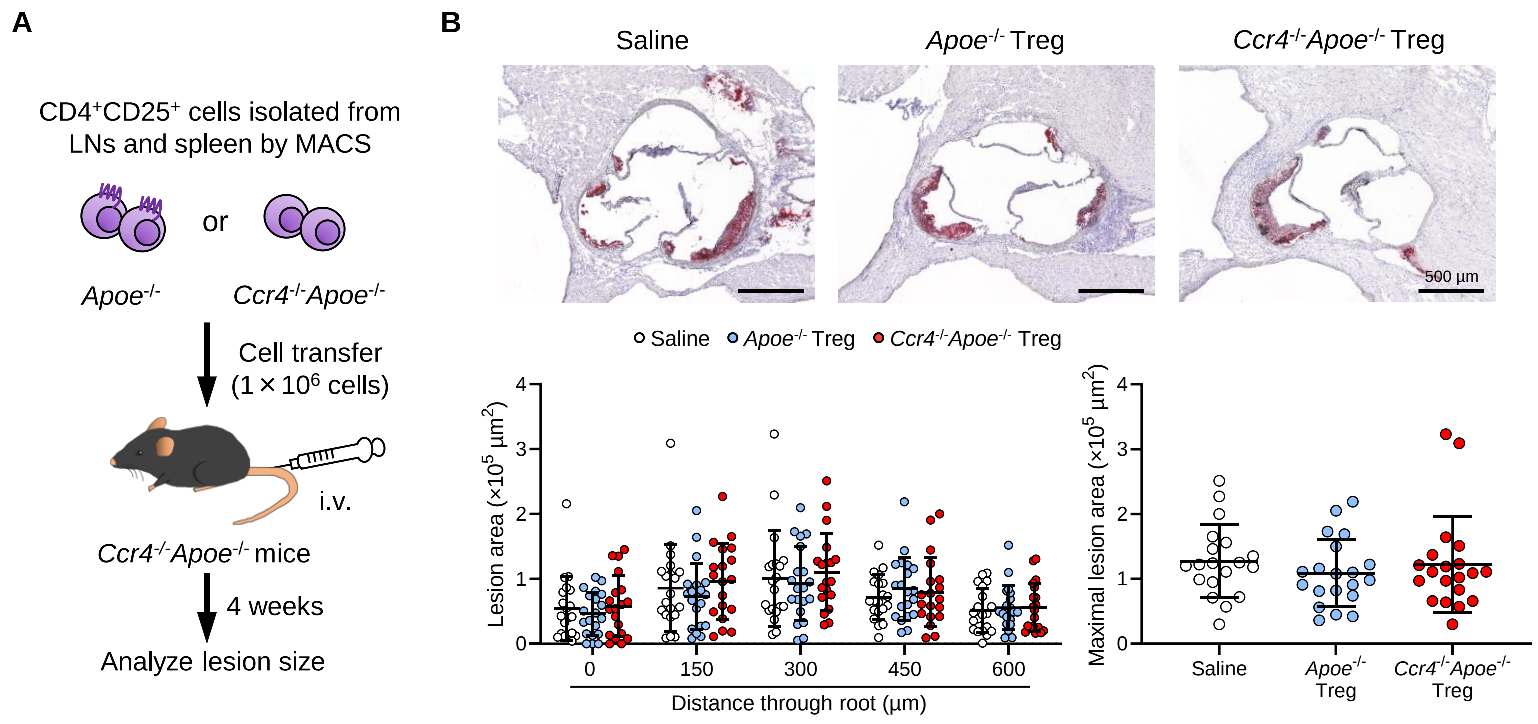


**Supplementary Figure 10. Transfer of CCR4-intact Tregs does not affect the development of early atherosclerotic lesions in *Ccr4$^{-/-}$Apoe$^{-/-}$* mice.**

**A**, Tregs purified from the peripheral LNs and spleen of *Apoe$^{-/-}$* or *Ccr4$^{-/-}$Apoe$^{-/-}$* mice were intravenously transferred into 12-week-old *Ccr4$^{-/-}$Apoe$^{-/-}$* mice fed a standard chow diet, and atherosclerotic lesions were analyzed at 16 weeks of age. As a control without cell transfer, 12-week-old *Ccr4$^{-/-}$Apoe$^{-/-}$* mice were intravenously injected with saline and atherosclerotic lesions were analyzed at 16 weeks of age. **B**, Representative photomicrographs of Oil Red O staining and quantitative analysis of atherosclerotic lesion area at 5 different levels and maximal lesions in the aortic sinus of *Ccr4$^{-/-}$Apoe$^{-/-}$* mice injected with saline (n=19), *Apoe$^{-/-}$* Tregs (n=20), or *Ccr4$^{-/-}$Apoe$^{-/-}$* Tregs (n=20). Black bars represent 500 µm as described. Data points represent individual animals. Horizontal bars represent means. Error bars indicate s.d.

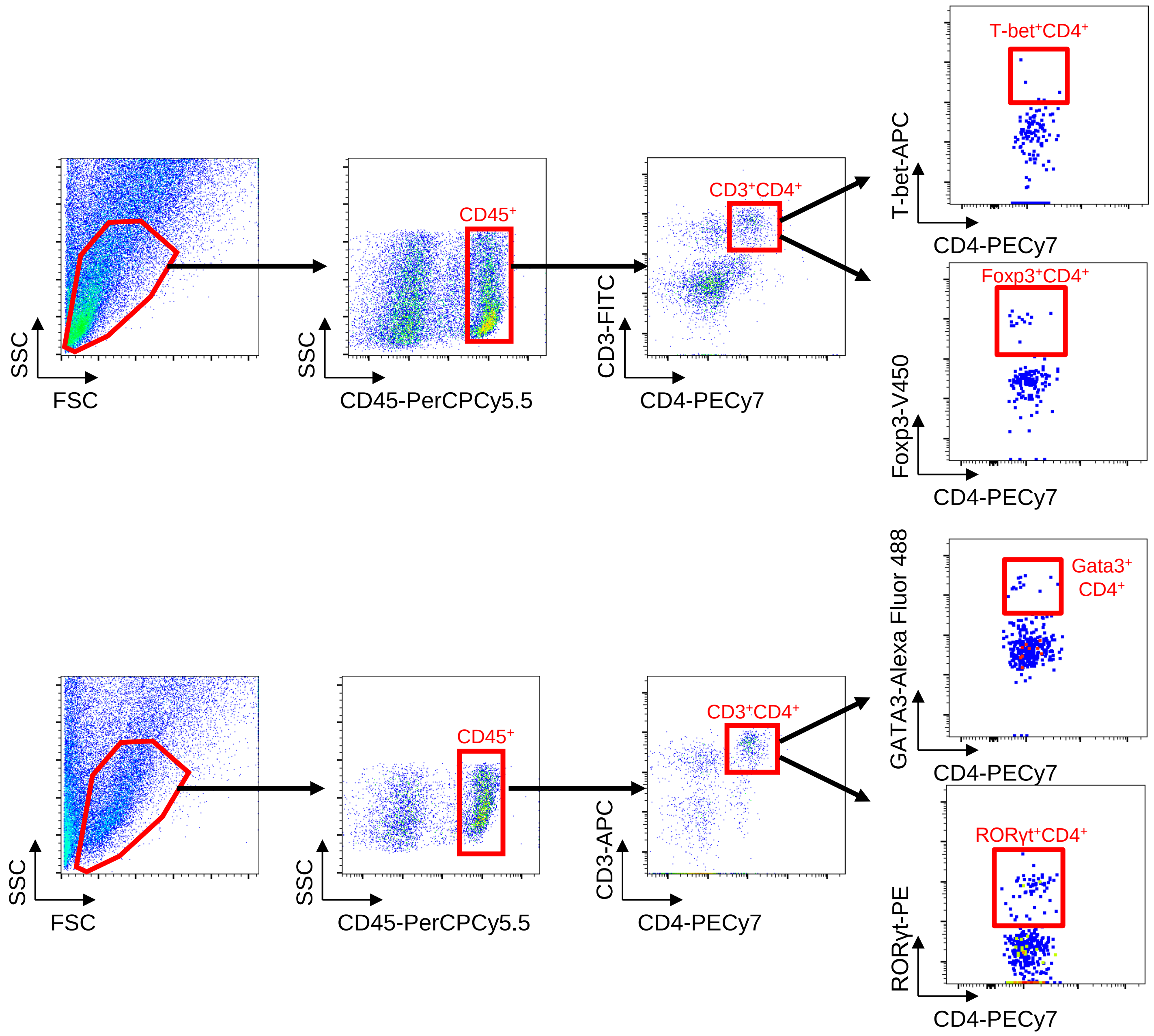


**Supplementary Figure 11. Gating strategy of flow cytometric analysis of T-bet, GATA3, RORγt, and Foxp3 expression in aortic $CD3^{+}CD4^{+}CD45^{+}$ T cells.**

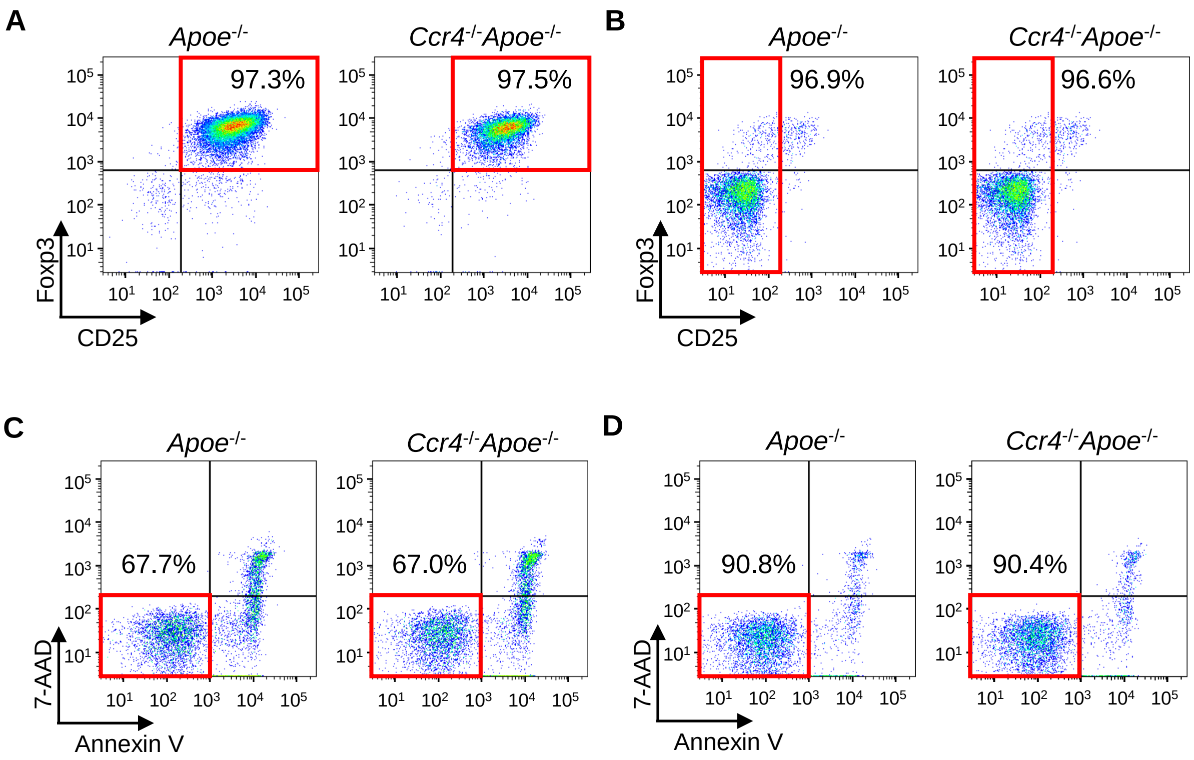


**Supplementary Figure 12. The purity and viability of CD4$^+$CD25$^+$ Tregs and CD4$^+$CD25$^-$ T cells isolated from *Apoe$^{-/-}$* or *Ccr4$^{-/-}$Apoe$^{-/-}$* mice.**

Representative flow cytometric analysis of CD25 and Foxp3 expression in CD4$^+$CD25$^+$ Tregs (**A**) and CD4$^+$CD25$^-$ T cells (**B**) purified from peripheral lymphoid tissues of *Apoe$^{-/-}$* or *Ccr4$^{-/-}$Apoe$^{-/-}$* mice. Representative flow cytometric analysis of the viability of CD4$^+$CD25$^+$ Tregs (**C**) and CD4$^+$CD25$^-$ T cells (**D**) purified from peripheral lymphoid tissues of *Apoe$^{-/-}$* or *Ccr4$^{-/-}$Apoe$^{-/-}$* mice. T cells which neither expressed 7-AAD nor Annexin V were considered viable.

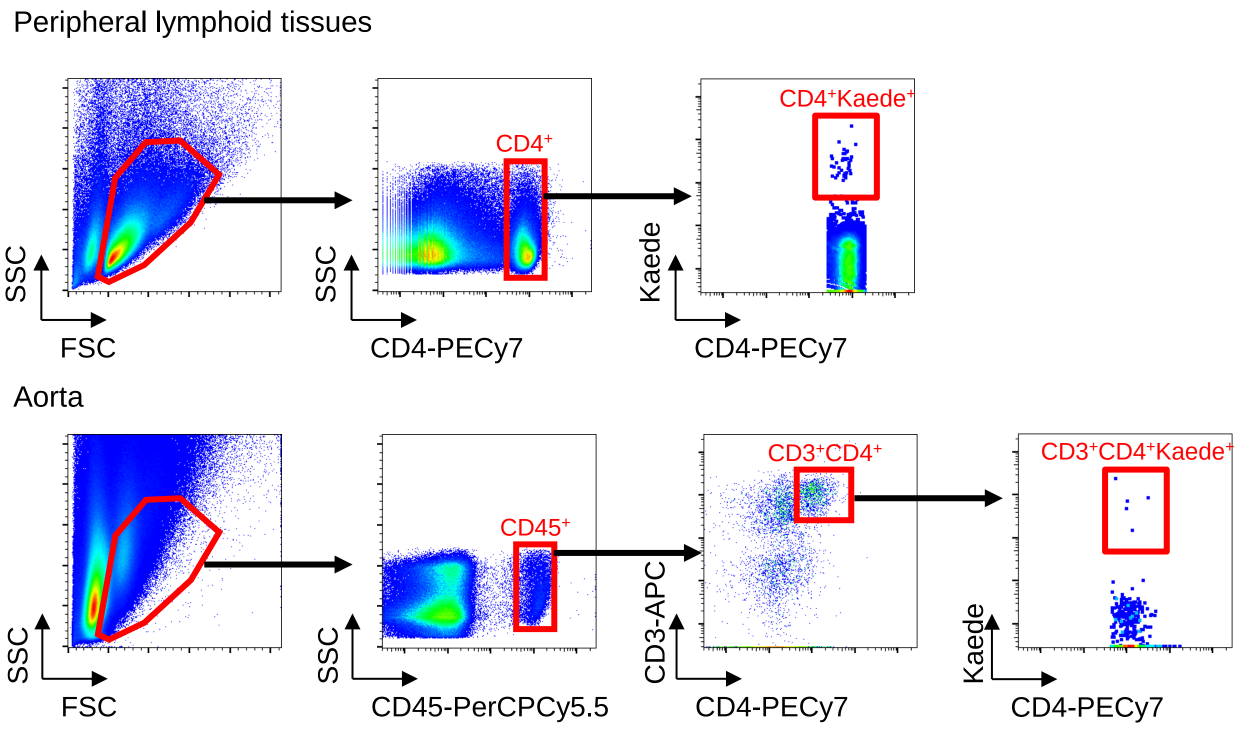


**Supplementary Figure 13. Gating strategy of flow cytometric analysis of Kaede-expressing Tregs in peripheral lymphoid tissues and aortas.**

#### Supplementary Tables

**Supplementary Table 1** Body weight and plasma lipid profile in 18-week-old mice

| | *Apoe*$^{-/-}$ | *Ccr4*$^{-/-}$*Apoe*$^{-/-}$ |
|---|---|---|
| Body weight (g) | 31.76 ± 1.94 (n=27) | 31.69 ± 2.24 (n=27) |
| Total cholesterol (mg/dL) | 526.5 ± 146.3 (n=10) | 520.4 ± 150.6 (n=10) |
| HDL-cholestetol (mg/dL) | 24.90 ± 6.49 (n=10) | 19.80 ± 5.87 (n=10) |
| Triglycerides (mg/dL) | 89.70 ± 21.07 (n=10) | 86.40 ± 39.58 (n=10) |

HDL, high-density lipoprotein.

**Supplementary Table 2**

Body weight and plasma lipid profile in *Apoe*$^{-/-}$ mice treated with saline, *Apoe*$^{-/-}$ Tregs, or *Ccr4*$^{-/-}$*Apoe*$^{-/-}$ Tregs

| | Saline | *Apoe*$^{-/-}$Treg | *Ccr4*$^{-/-}$*Apoe*$^{-/-}$Treg |
|---|---|---|---|
| Body weight (g) | 27.82 ± 2.82 (n=18) | 27.80 ± 2.42 (n=20) | 27.72 ± 2.59 (n=19) |
| Total cholesterol (mg/dL) | 505.3 ± 114.0 (n=10) | 605.5 ± 105.1 (n=10) | 607.3 ± 76.35 (n=10) |
| HDL-cholestetol (mg/dL) | 15.00 ± 2.87 (n=10) | 15.20 ± 3.26 (n=10) | 13.80 ± 4.83 (n=10) |
| Triglycerides (mg/dL) | 87.40 ± 36.69 (n=10) | 92.70 ± 27.15 (n=10) | 96.60 ± 33.94 (n=10) |

HDL, high-density lipoprotein.

**Supplementary Table 3**

Body weight and plasma lipid profile in *Ccr4*$^{-/-}$*Apoe*$^{-/-}$ mice treated with saline, *Apoe*$^{-/-}$ Tregs, or *Ccr4*$^{-/-}$*Apoe*$^{-/-}$ Tregs

| | Saline | *Apoe*$^{-/-}$Treg | *Ccr4*$^{-/-}$*Apoe*$^{-/-}$Treg |
|---|---|---|---|
| Body weight (g) | 32.44 ± 2.75 (n=12) | 30.69 ± 3.55 (n=12) | 32.16 ± 2.12 (n=12) |
| Total cholesterol (mg/dL) | 589.1 ± 144.3 (n=10) | 616.1 ± 105.6 (n=10) | 526.4 ± 115.9 (n=10) |
| HDL-cholestetol (mg/dL) | 16.30 ± 3.65 (n=10) | 13.80 ± 3.19 (n=10) | 12.10 ± 2.96* (n=10) |
| Triglycerides (mg/dL) | 83.0 ± 57.56 (n=10) | 91.60 ± 55.52 (n=10) | 72.70 ± 13.99 (n=10) |

HDL, high-density lipoprotein. *$P$<0.05 vs. saline-treated mice; 1-way ANOVA followed by Tukey's multiple comparisons test.

**Supplementary Table 4** Antibodies for immunohistochemistry

| Antibodies | Clone | Conjugate | Source |
|---|---|---|---|
| anti-CCL17 Ab | - | - | abcam<br>ab182793 |
| anti-MOMA-2 Ab | - | - | BMA Biomedical<br>T-2007 |
| anti-CCL22 Ab | - | - | R&D<br>AF439 |
| anti-CD4 Ab | RM4-5 | - | BD Biosciences<br>550280 |
| anti-Rabbit IgG | - | Alexa Fluor 568 | Thermo Fisher Scientific<br>A11011 |
| anti-Goat IgG | - | Alexa Fluor 568 | Thermo Fisher Scientific<br>A11057 |
| anti-Rat IgG | - | Alexa Fluor 488 | Thermo Fisher Scientific<br>A21208 |
| anti-Rat IgG | - | Biotin | abcam<br>ab102250 |

**Supplementary Table 5** Antibodies for flow cytometry

| Antibodies | Clone | Fluorescent dye | Source |
|---|---|---|---|
| anti-CD16/CD32 Ab | 2.4G2 | - | BD Pharmingen 553142 |
| anti-CD45 Ab | 30-F11 | PerCPCy5.5 | BD Pharmingen 550994 |
| anti-CD3 Ab | 145-2C11 | | BD Pharmingen |
| | | APC | 553066 |
| | | FITC | 553062 |
| | | PECy7 | 552774 |
| anti-CD4 Ab | RM4-5 | PECy7 | BD Pharmingen 552775 |
| anti-CCR4 Ab | 2G12 | PE | BioLegend 131204 |
| anti-Foxp3 Ab | FJK-16s | | eBioscience |
| | | V450 | 48-5773-82 |
| | | APC | 17-5773-82 |
| anti-CD44 Ab | IM7 | PE | BD Pharmingen 553134 |
| anti-CD62L Ab | MEL-14 | FITC | BD Pharmingen 553150 |
| anti-Ki-67 Ab | SolA15 | FITC | eBioscience 11-5698-82 |
| anti-CD152 Ab | UC10-4B9 | APC | eBioscience 17-1522-82 |
| anti-CD103 Ab | M290 | FITC | BD Pharmingen 557494 |
| anti-CD25 Ab | PC61 | PE | BD Pharmingen 553866 |
| anti-IFN-γ Ab | XMG1.2 | PE | BD Pharmingen 554412 |
| anti-IL-4 Ab | 11B11 | PE | BD Pharmingen 554435 |
| anti-IL-10 Ab | JES5-16E3 | APC | BD Pharmingen 554468 |
| anti-IL-17 Ab | TC11-18H10 | APC | BD Pharmingen 560184 |
| anti-T-bet Ab | 4B10 | APC | BioLegend 644814 |
| anti-Gata3 Ab | L50-823 | Alexa Fluor 488 | BD Pharmingen 560163 |
| anti-RORγt Ab | Q31-378 | PE | BD Pharmingen 562607 |
| anti-CD8 Ab | 53-6.7 | PerCPCy5.5 | BD Pharmingen 553033 |
| anti-B220 Ab | RA3-6B2 | PE | BD Pharmingen 553090 |
| anti-CD11b Ab | M1/70 | V450 | BD Horizon 560455 |
| anti-Ly6G Ab | 1A8 | FITC | BD Pharmingen 551460 |
| anti-Ly6C Ab | AL-21 | APC | BD Pharmingen 560595 |
| anti-NK1.1 Ab | PK136 | APC | BD Pharmingen 550627 |
| anti-CD11c Ab | HL3 | V450 | BD Horizon 560521 |
| anti-I-Ab Ab | AF6-120.1 | FITC | BD Pharmingen 553551 |
| anti-CD80 Ab | 16-10A1 | PE | BD Pharmingen 553769 |
| anti-CD86 Ab | GL1 | APC | BD Pharmingen 558703 |
| anti-CD279 Ab | 29F.1A12 | FITC | BioLegend 135213 |

**Supplementary Table 6** Primer sequences for quantitative reverse transcription PCR

| Gene | Forward primer sequence （5'→3'） | Reverse primer sequence （5'→3'） |
|---|---|---|
| *Gapdh* | TGTGTCCGTCGTGGATCTGA | TTGCTGTTGAAGTCGCAGGAG |
| *Il1b* | TCCAGGATGAGGACATGAGCAC | GAACGTCACACACCAGCAGGTTA |
| *Il6* | CCACTTCACAAGTCGGAGGCTTA | GCAAGTGCATCATCGTTGTTCATAC |
| *Il10* | GACCAGCTGGACAACATACTGCTAA | GATAAGGCTTGGCAACCCAAGTAA |
| *Tnf* | CCACCACGCTCTTCTGTCTAC | AGGGTCTGGGCCATAGAACT |
| *Ifng* | CGGCACAGTCATTGAAAGCCTA | GTTGCTGATGGCCTGATTGTC |
| *Tbx21* | CTGCCTACCAGAACGCAGA | AAACGGCTGGGAACAGGA |
| *Gata3* | GGATGTAAGTCGAGGCCCAAG | ATTGCAAAGGTAGTGCCCGGTA |
| *Rorc* | CACAGAGACACCACCGGACAT | CGTGCAGGAGTAGGCCACATT |
| *Foxp3* | CTCATGATAGTGCCTGTGTCCTCAA | AGGGCCAGCATAGGTGCAAG |
| *Ctla4* | CCTCTGCAAGGTGGAACTCATGTA | AGCTAACTGCGACAAGGATCCAA |
| *Cd103* | ATGGCATTCAGTGGTCTGTGCTA | CACCAAGGATCGGCAGTTCA |
| *Tnfrsf18* | GTTCAGAACGGAAGTGGCAACA | GCTTGCAGATCTTGCACTGAGG |
| *Tgfb* | GTGTGGAGCAACATGTGGAACTCTA | TTGGTTCAGCCACTGCCGTA |
| *Cd44* | CTGGCACTGGCTCTGATTCTTG | TCCCATTGCCACCGTTGA |
| *Cd69* | TGGCCCAACGCTCTTGTTC | GCCCAATCCAATGTTCCAGTTC |
| *Ccr4* | TCTACAGCGGCATCTTCTTCAT | CAGTACGTGTGGTTGTGCTCTG |
| *Ccr5* | CCTAGCCAGAGGAGGTGAGACATC | AGCTATAGGTCGGAACTGACCCTTG |
| *Ccr6* | GGCAGTTACTCATGCCACCAA | GGAGCAGCATCCCACAGTTAAAG |
| *Ccr7* | GGTGGTGGCTCTCCTTGTCATT | ACACCGACTCGTACAGGGTGTAGTC |
| *Ccr8* | CAGACCCACAACCTGCTGGA | GACAGCGTGGACAATAGCCAGA |
| *Ccr1* | GGTTGGGACCTTGAACCTTG | GGGTAGGCTTCTGTGAAATCTG |
| *Cxcr3* | ATCACCTGGTGGTGCTAGTGGA | AAAGGCATAGAGCAGCGGATTG |
| *Cx3cr1* | AAGCACTTGCCTCTGGTGGA | AGGCCTCAGCAGAATCGTCATA |
